# S494 O-glycosylation site on the SARS-COV-2 RBD Affects the Virus Affinity to ACE2 and its Infectivity; A Molecular Dynamics Study

**DOI:** 10.1101/2020.09.12.294504

**Authors:** Shadi Rahnama, Maryam Azimzadeh Irani, Mehriar Amininasab, Mohammad Reza Ejtehadi

**Author notes:** O-glycosylation of the SARS-COV-2 RBD.

## Abstract

SARS-COV-2 is a strain of Coronavirus family which caused the extensive pandemic of COVID-19, which is still going on. Several studies showed that the glycosylation of virus spike (S) protein and the Angiotensin-Converting Enzyme 2 (ACE2) receptor on the host cell is critical for the virus infectivity. Molecular Dynamics (MD) simulations were used to explore the role of a novel mutated O-glycosylation site (D494S) on the Receptor Binding Domain (RBD) of S protein. This site was suggested as a key mediator of virus-host interaction. We showed that the decoration of S494 with elongated O-glycans results in stabilized interactions on the direct RBD-ACE2 interface with more favorable binding free energies for longer oligosaccharides. Hence, this crucial factor must be taken into account for any further inhibitory approaches towards RBD-ACE2 interaction.

## Introduction

Severe Acute Respiratory Syndrome Corona Virus 2 (SARS-COV-2) is a positive-sense single-stranded RNA virus which caused the pandemic of COVID-2019 that is still going on. In Wuhan, China, the virus is believed to transmit from bat to human^1^ and underwent several inter and intra species passages. This large-scaled pandemic motivated several scientific groups to control the virus’s spread and treat the infected patients by all means. The researchers are mainly focused on the phylogeny and origin of SARS-COV-2,^1^ Structural assembly and the dynamics of virion-host cell proteins^2–4^ and the experimental efforts to obtain a SARS-COV-2 neutralizing antibody.^5,6^

The SARS-COV-2 virion assembly and its infusion into the human cell are similar to the one known for SARS-COV.^2^ The fusion protein known as the spike (S) protein interacts with ACE2 on the host cell via its Receptor Binding Domain (RBD)^3,7^ (Figure 1).The RBD-ACE2 interaction takes place via four *β* strands on RBD (*β*4-*β*7) and two *α* helices on ACE2 binding interface (H1,H2)^3^(Figure 1). Electron microscopy and X-ray crystal structures elucidated the active form of SARS-COV-2 S protein which assembles as a trimer,^2^ and one RBD of one S protein complex ACE2, ^3^ respectively. The structures of S protein in the apo and ACE2 bound complexes show that RBD adopts a 3D fold similar to the RBD of SARS-COV;.^2^ However, six novel mutations on SARS-COV-2 RBD could lead to different binding affinity to ACE2. ^1^ Speculations regarding SARS-COV-2-ACE2 binding affinity suggested two different scenarios. One reported a dramatic increase (up to 20-fold) in the binding affinity of SARS-COV-2 to ACE2 compared to SARS-COV.^2^ While, the other suggests a similar binding affinity for SARS-COV-2 and SARS-COV to ACE2.^3,8^ Either way, the mutations in SARS-COV2 RBD certainly results in attenuated binding affinity of all designed inhibitors which successfully work on SARS-COV.^2^ It was a challenging issue for anti-SARS-COV-2 drug design attempts since the pandemic has started.

**Figure 1:**
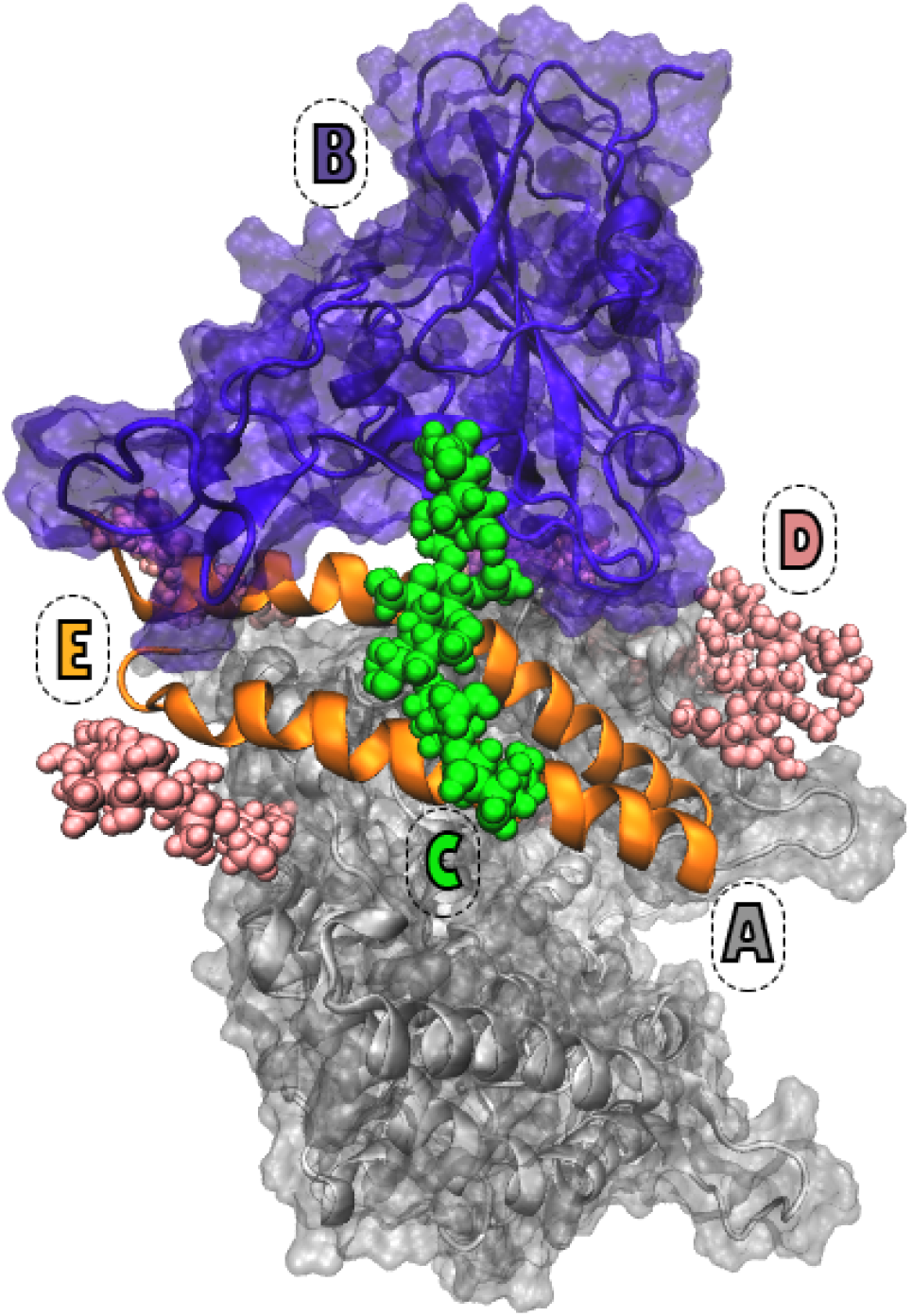
The extracellular domain of human ACE2 and RBD of SARS-COV-2 S protein is shown with gray and purple, respectively (A and B). The RBD attached O-glycan is shown with green spheres (C). For the sake of simplicity, only the N-glycans in the proximity of RBD-ACE2 interface are shown with pink spheres. Visualization of the complex with full presentation of the N-glycans can be found in Figure 11 (D). Two O-glycan interaction sites H1 and H2, are shown with orange (E).

In addition to the 3D structures of RBD-ACE2 complex, several recent Molecular Dynamics (MD) studies addressed the complex dynamics to design peptide inhibitors. That block the ACE2-RBD interaction and evaluate the currently targeted binding epitopes’ accessibility for further improvements. ^4,9,10^ Earlier MD studies explored the effect of pH ^11^ and temperature^12^ on the dynamics of SARS-COV-ACE2 complex.To the best of our knowledge, there is currently no approved therapeutic inhibitor for SARS-COV-2.

A pioneering study showed that the observed mutations at the junction of S1 and S2 subdomains of the SARS-COV-2 S protein result in the emergence of a polybasic cleavage site adjacent to three O-glycosylation sites which are novel to SARS-COV-2 and were not observed in any other related virion.^1^ It was proposed that O-glycosylation (addition of glycan building blocks to hydroxyl oxygen of Ser/Thr residues which is an enzymatic post transnational modification^13^) could lead to recognition of polybasic cleavage site by Furin enzyme and thus resulted in higher infectivity and broadens the host range of the virus.^1^

A Recent comprehensive experimental mass-spectrometry study on both N- and O-glycans attached to the spike protein did not report O-glycosylation of the S494. However, two O-glycosylated sites (S325/T323) were reported on the RBD.^14^ Another recent computational study that was validated by experimental data reported that the S325/T323 O-glycosylation sites that were observed by Shajahan et al. is present with 11% frequency along side other O-glycosylation site that were decorated with lower frequencies. ^15^

Six mutations have been reported on the RBD of SARS-COV-2 compared to the SARS-COV. One of these six occurring mutations is D494S on the RBD of S protein. Based on the proposed mechanism for the O-glycosylation of Furin cleavage site and the Evidences mentioned above,^1,14,15^ we propose that Serine494 of SARS-COV-2 RBD could also become O-glycosylated even though it is not experimentally confirmed (Figure 1).

Our comprehensive review of literature on pathology and glycosylation pattern of the envelope proteins of all other coronaviruses and their close relatives showed strong evidence for the critical role of O-glycosylation in regulation and immune evasion of the viruses. ^1,16–18^ Also, former studies showed that the O-glycosylation of the membrane (M) protein of Mouse Hepatitis Virus (MHV) triggers a lower interferon level than the N-glycosylated M protein.^16^ 3A protein of the MHV, an accessory protein to the M protein is also known to be O-glycosylated. Thus, it could be shielded from Peptide-N Glycosidase F (PNGase F) which cleaves most monosaccharide attached to protein. On the other hand, a complex non-template mechanism of the O-glycosylation could occur via several Polypeptide N-acetyl Galactoseamine Transferases (PPGalNAcTs) specific to various tissues was shown to be a common mechanism for the pathology of several high-evasion viruses such as Influenza. ^19^ In fact, the experimental findings suggest that the mucin proteins of human respiratory airways are heavily O-glycosylated with carbohydrate chains.^20–22^ These dense O-glycan chains are known to trap invading microorganisms such as various strains of bacteria.^23^

Putting all the information mentioned above together, it is highly plausible that D494S mutation has an evolutionary origin and leads to O-glycosylation of RBD in the SARS-COV-2. Hence, the O-glycosylation could be the overlooked factor for the SARS-COV-2 increased binding affinity to ACE2, explaining the virus’s high infectivity in the human host and exposing people with a higher level of blood sugar at higher risk. ^24^ Our observations may explain the reason for the higher vulnerability of diabetic patients to infection.

Herein, we explored this possibility by modeling the O-glycosylated RBD structure and dynamics in interaction with ACE2 (Figure 2). We show that O-glycosylation does indeed increase SARS-COV-2 RBD-ACE2 binding affinity. Also, we show that attachment of the elongated O-glycans will increase the binding affinity. Furthermore, we also considered the structural role of N-glycosylation (addition of glycan building blocks to the nitrogen atom of Asn located at the Asn-X-Ser/Thr motif, enzymatic post-translationalmodification) of both RBD and ACE2 in all of the simulations.

**Figure 2:**
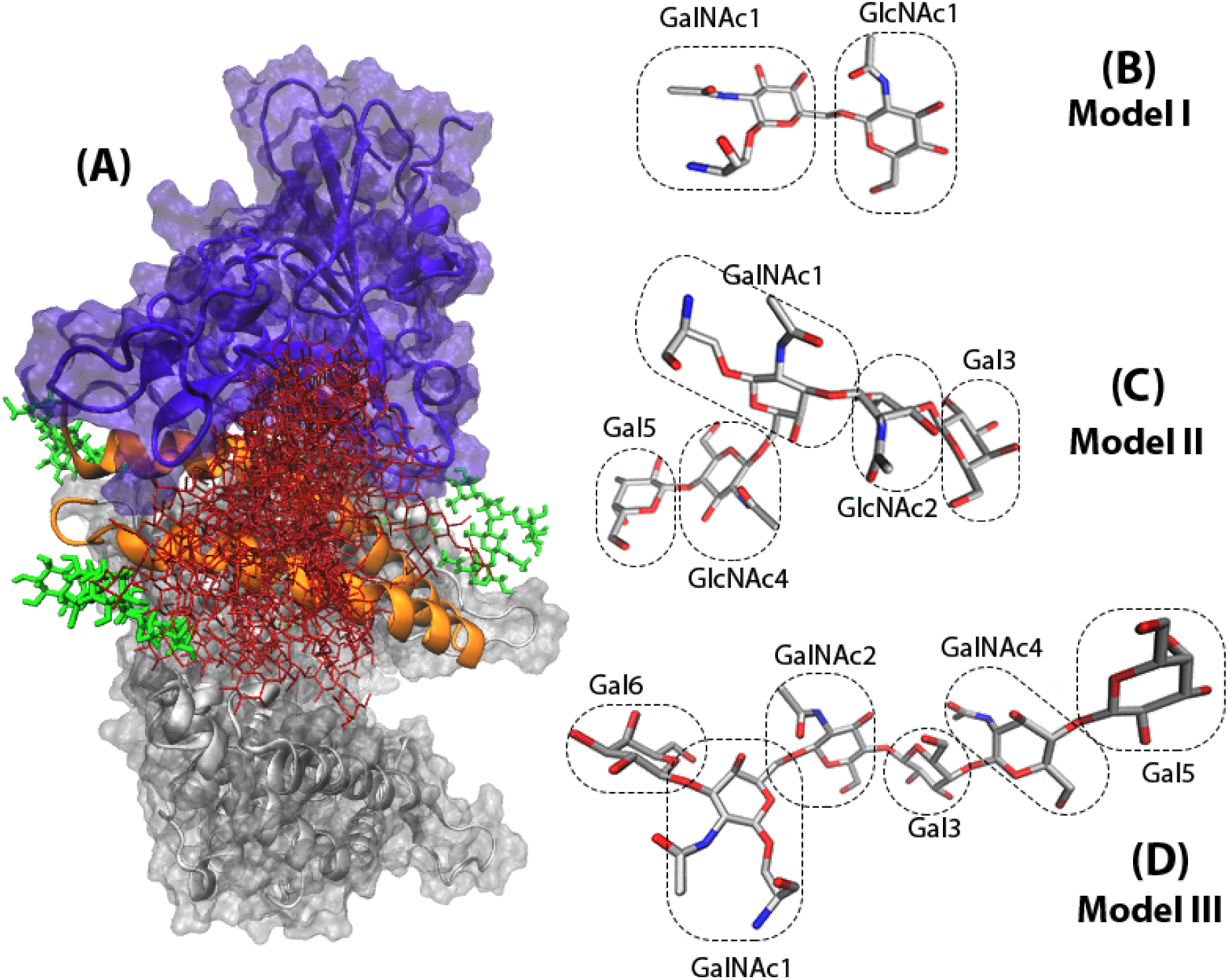
(A) The overlaid presentation of O-glycosylated RBD-ACE2 complex is shown with red sticks. The extracellular domain of human ACE2 and RBD of SARS-COV-2 S protein is shown with gray and purple, respectively. For the sake of simplicity, only the N-glycans in the proximity of RBD-ACE2 interface are shown with green sticks. Visualization of the complex with full presentation of the N-glycans can be found in Figure 11. O-glycan models I (B), II (C) and III (D) are shown with atom name sticks.

In summary, this work provides strong evidence that O-glycosylation of SARS-COV2 RBD in the respiratory airways leads to stronger interactions between the virion and the host cell receptor. Hence, further inhibitor design attempts must take the detailed atomistic glycan-protein interaction reported here into account as a critical factor for future investigations.

## Methods

### O- and N-glycosylated models of RBD-ACE2 complexes and Molecular Dynamics simulations

The crystal structure of SARS-COV-2 RBD (residue 333 to 526) in complex with the extracellular domain of ACE2 (residue 19 to 615) (PDB ID: 6M0J) was used as the starting structure. Zn^2+^ and Cl^−^ ions which were resolved in the crystal structure for their crucial role in stabilizing ACE2 structure were also preserved at S1 and S2 subunits of ACE2. All Aspargines located within N-glycosylation motifs from RBD (Asn343) and ACE2 (Asn53, Asn90, Asn103, Asn322, Asn432, and Asn546) were glycosylated. The GLYCAM online builder^25^ was used to attach the oligosaccharide glycans to each site. This oligosaccharide model was selected as a primary model of glycosylation which occurs in normal human cells.^26^ Supporting table 1 provides details on the structure of attached N-glycans.

O-glycosylation of Serine494 of RBD was carried out by attaching three models of core and branched typical human O-glycans using GLYCAM online builder.^25^ Models I, II, and III are consist of two, five, and six monosaccharide units, respectively (Figure 2). These O-glycan models were attached to RBD, while RBD-ACE2 complex was kept fully N-glycosylated (Figure 1). The RBD-ACE2 model without attached O-glycan will be referred to as model 0 in the rest of the text and figures. All systems were solvated using TIP3P water model^27^ and were neutralized by attaining a buffered environment in 150 mM NaCl. The resulting water box approximately has a dimension of 121 × 111 × 149 Å^3^ and are typically consist of 190000 atoms each. All systems were parameterized with the CHARMM36m force field,^28,29^ and NAMD2.12^30^ was used to perform the MD simulations. Periodic boundary conditions under NPT ensemble was applied, while employing Langevin thermostat and Nose–Hoover Langevin piston method to keep systems’ temperature and pressure at 310 K and 1 bar, respectively. A cutoff of Å was assigned for computation of short range nonbonded van der Waals interactions. The particle mesh Ewald method^31^ was used for long-range electrostatic interactions. R-RESPA multiple time-step schemes were used for the integration of motion equations. ^30^ Using this integrator, the Lennard-Jones interactions and bonded ones were updated every step and electrostatic interactions every two steps. Along with the restraining of all covalent hydrogen bonds by SHAKE, the time steps for integration were set to 2fs. Final models of all complexes were minimized for 5000 steps to remove any steric clashes. Following minimization, the systems firstly equilibrated in NVT ensemble carried out for 0.5 ns. Subsequently, system equilibration was continued in NPT ensemble for an additional 0.5 ns. During both steps, the position of atoms in the complex (proteins, Zn^+2^ and Cl^−1^ ions) was restrained using a harmonic potential with spring constant k = 1 kcal/(molÅ^2^). Production runs were then performed in three replicates of each system for 50ns. During the production runs, all restraints on the protein were turned off, while dummy bonds between Zn^+2^ and Cl^−1^ ions and their adjunct atoms within radius of 3 Å were kept intact.

### Binding free energy calculations

The binding free energy for all sets of ligand-receptor complexes was calculated with Molecular Mechanics Poison Boltzmann Surface Area (MMPBSA). Here we used CaFE pipeline tool^32^ on 250 frames from the last 10 ns of MD simulations and an ensemble-averaged overall replica of each model. For electrostatic energy calculations, APBS method ^33^ was used. The dielectric constants for solvent and protein were set to 80.0 and 1.2 respectively (More details could be found in Supplementary method).Probe radius for calculation of SASA was set to 1.4 Å.

### MD simulations analyses

Root Mean Squared Deviation (RMSDs), Root Mean Squared Fluctuation (RMSFs), and the distances between RBD and ACE2, H1 and H2 were calculated by tcl scripts taking advantage of VMD.^34^ RMSD of each system was calculated by considering the backbone atoms (C_*α*_, C, and N). RMSF and the distance among the geometric centers of domains were calculated for C_*α*_ atoms. All trajectories were fitted to the starting conformation to present the conformational changes during simulation time. All the plots were generated by Matplotlib^35^ library of Python^36^ and the figures by VMD^34^ and PyMOL.^37^

## Results and Discussion

### Dynamics of the O- and N- glycosylated RBD-ACE2

It is worth mentioning again that as all the systems are fully N-glycosylated here, the N-glycans’ global effect is explicit in the dynamics. However, the interpretation of results is mainly focused on O-glycosylation.

In the rest of the text and figures, RMSD and RMSF values are averaged over the three replicate simulations of each model and the light shades around the plots presents the error bars of each average plot. The histograms at the right side of all RMSD and RMSF plots are calculated for the last 30 ns of the average plots.

RMSD plots of RBD-ACE2 complex simulations show that among three O-glycosylated systems, model III is the least flexible one, while models II and I come just after in order (Figure 3 A and B). Interestingly, Model 0 falls between models II and I (Figure 3). Thus, as it is expected, the length of attached O-glycan is important for RBD-ACE2 interactions. The longer glycans lead to a more stabilized RBD-ACE2 interactions. It could play a significant role in the high evasion of SARS-COV-2. The histograms of the Probability Density Function (PDF) for RMSD values show a clear decrease in the peaks of models III and II with less fluctuation amplitude compared to models I and 0 Figure 3.

**Figure 3:**
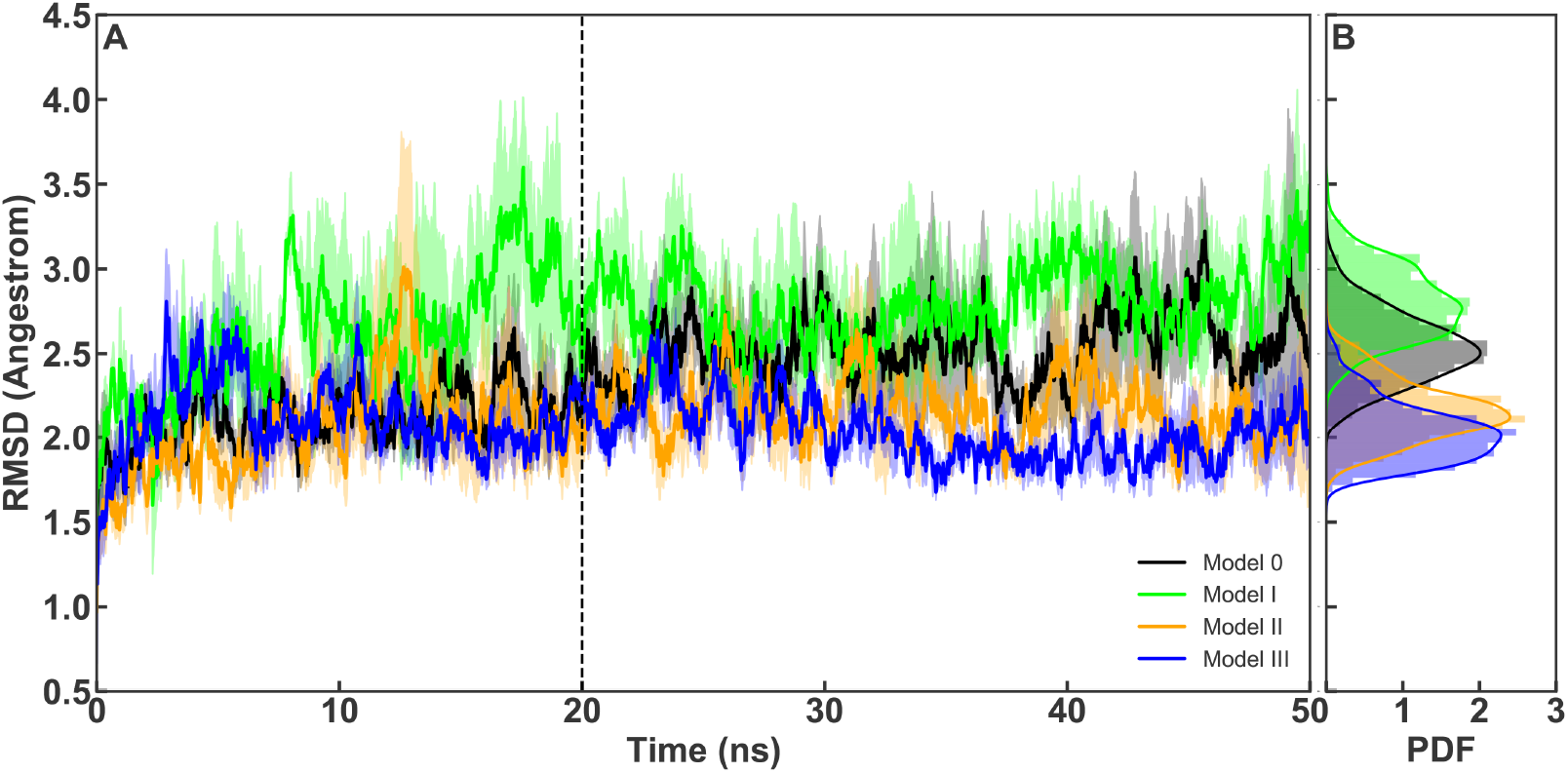
(A) Average backbone RMSD plots for RBD-ACE2, calculated from all replicates of each system and are shown for model 0 (black), model I (lime), model II (orange) and model III (blue). Light shades around each plot presents standard error for each calculation. (B) Probability Density Function (PDF) of RMSD sampled over the last 30 ns (dashed line) of the simulations are shown in histograms.

RMSF plots are also consistent to these observations (Figures 4,5) and show that the most dramatic increase of fluctuations occurs in residues 430 to 450 of RBD in model I. While, models II and III and the 0 present similar fluctuation patterns (Figure 4). Two residues (GLY446 and Tyr449) within this region are known to interact with the ACE2 directly.^38^ These results are interesting as we know which due to the experimental findings, the O-glycans of respiratory airways are often elongated oligosaccharides.^23^

**Figure 4:**
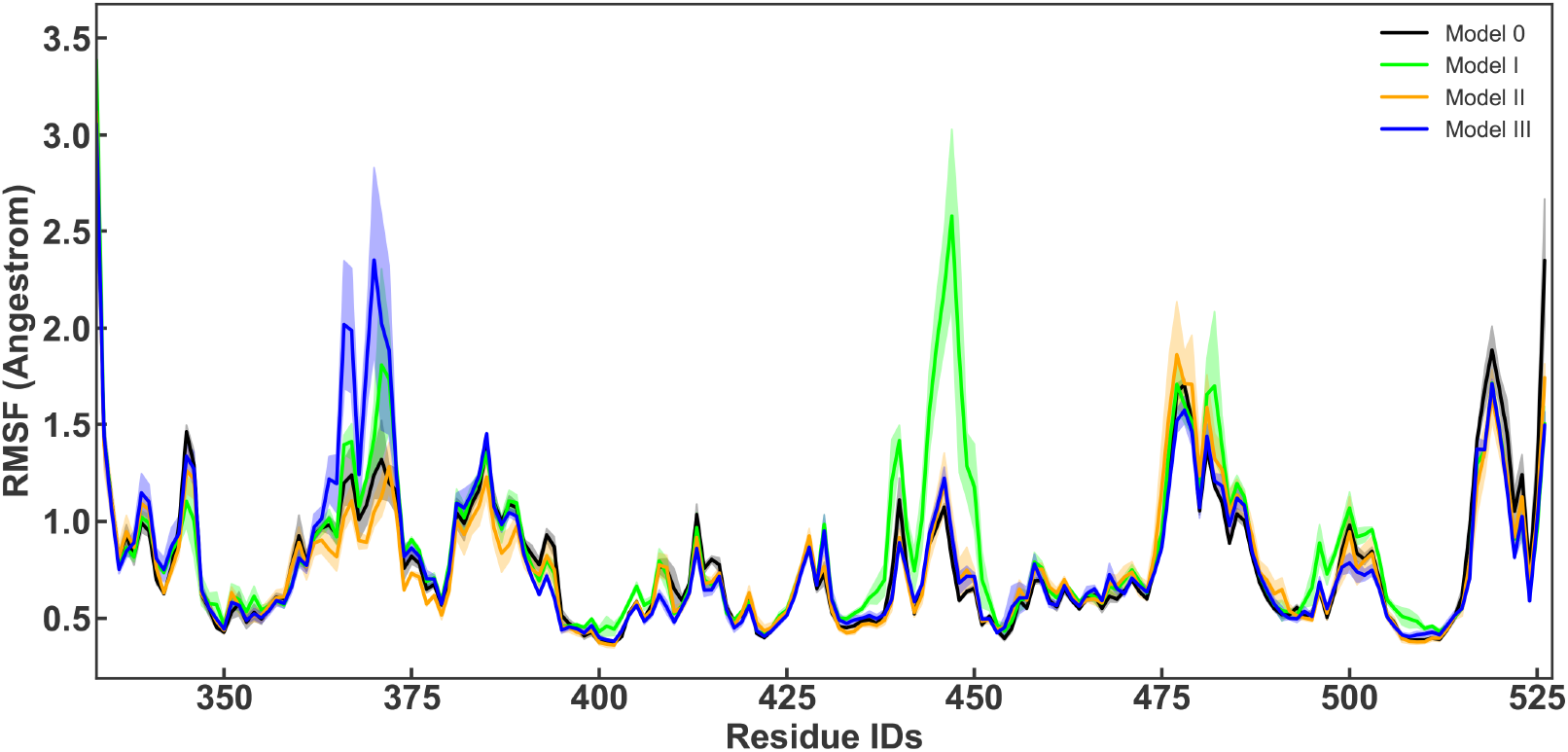
RMSF per residue of RBD calculated from all replicates of each system and are shown for model 0 (black), model I (lime), model II (orange) and model III (blue). Light shades around each plot presents standard error for each calculation.

**Figure 5:**
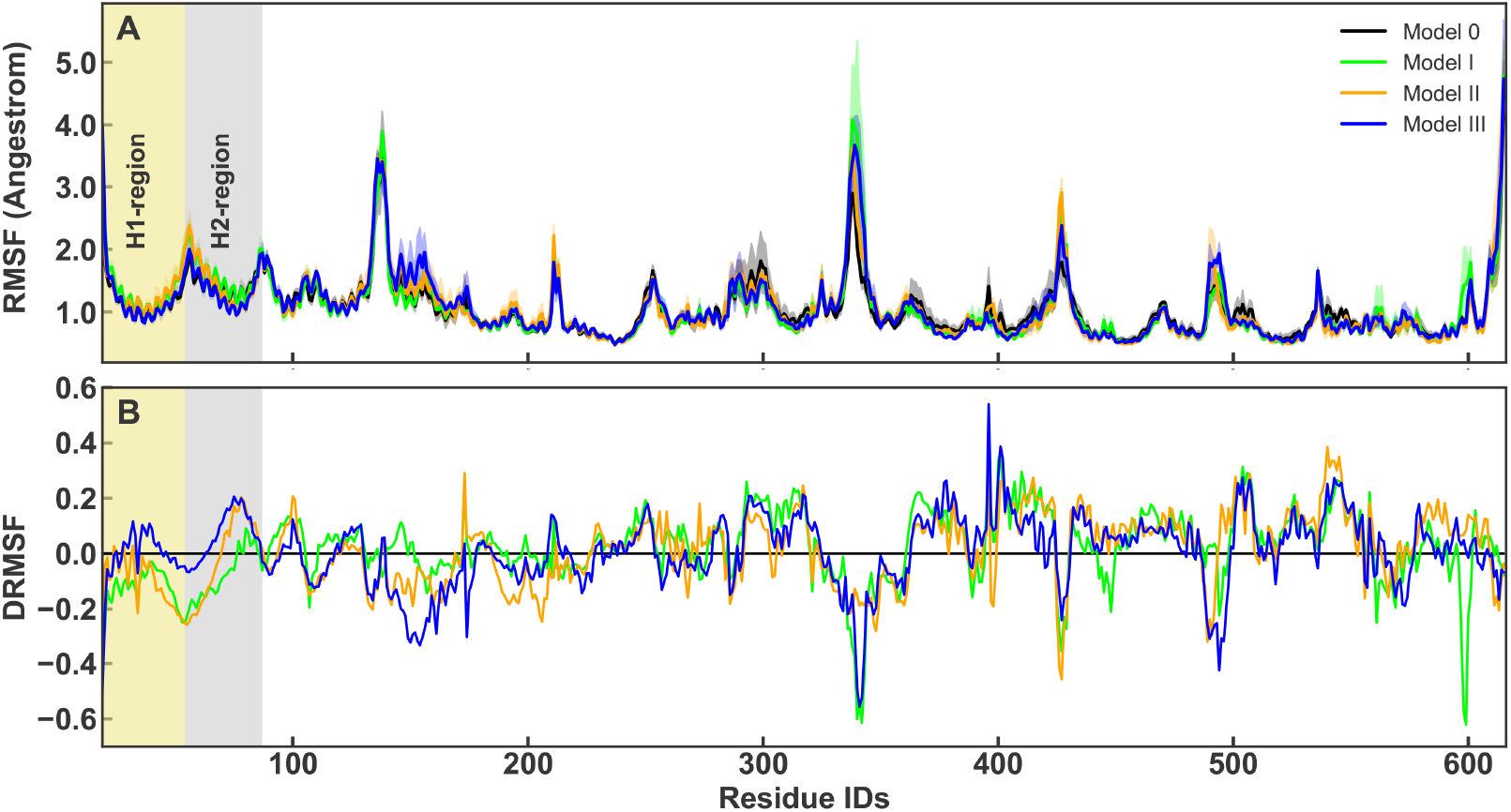
(A) RMSF per residue of ACE2 calculated from all replicates of each system and are shown for model 0 (black), model I (lime), model II (orange) and model III (blue). Light shades around each plot presents standard error for each calculation. The transparent boxes indicates H1 and H2. (B) The relative difference in RMSF, *D*_RMSF_ is shown per residue for all the models (with the same color codes).

### Detailed interactions

Detailed investigations show that decreased flexibility is more pronounced for RBD (Figure 6), while ACE2 average RMSD plots present a noticeable overlap among all systems (Figure 7.A-C).

**Figure 6:**
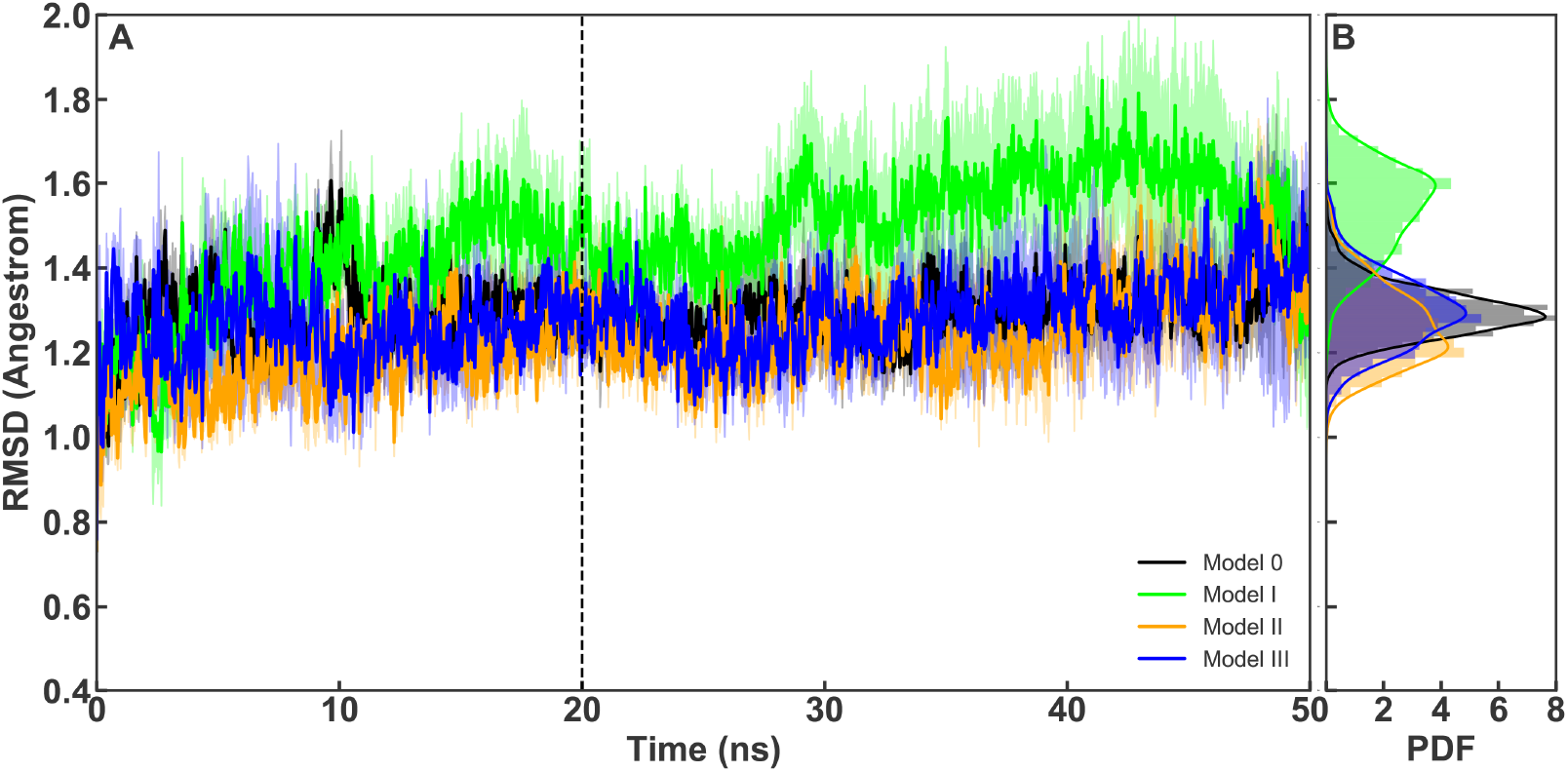
(A) Average backbone RMSD plots for Receptor binding RBD calculated from all replicates of each system and are shown for model 0 (black), model I (lime), model II (orange) and model III (blue). Light shades around each plot presents standard error for each calculation. (B) Probability Density Function (PDF) of RMSD sampled over the last 30 ns (dashed line) of the simulations are shown in histograms.

**Figure 7:**
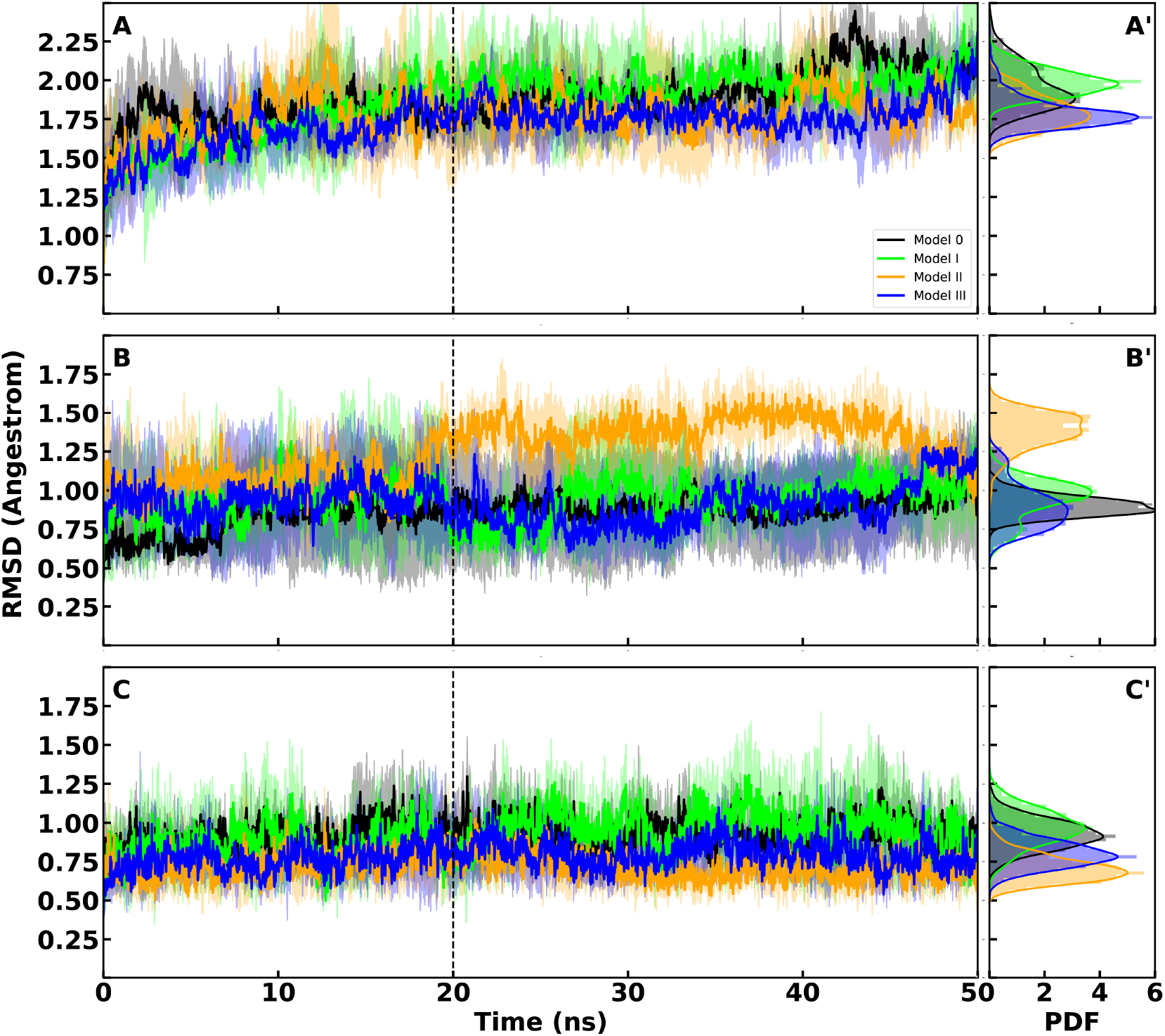
(A) Average backbone RMSD plots for ACE2 calculated from all replicates of each system and are shown for model 0 (black), model I (lime), model II (orange) and model III (blue). Light shades around each plot presents standard error for each calculation. (A’) Probability Density Function (PDF) of RMSD sampled over the last 30 ns (dashed line) of the simulations are shown in histograms. (B) Average backbone RMSD plots for *α*-Helix1 calculated from all replicates of each system and are shown for model 0 (black), model I (lime), model II (orange) and model III (blue). Light shades around each plot presents standard error for each calculation. (B’) Probability Density Function (PDF) of RMSD sampled over the last 30ns (dashed line) of the simulations are shown in histograms.(C) Average backbone RMSD plots for *α*-Helix2 calculated from all replicates of each system and are shown for model 0 (black), model I (lime), model II (orange) and model III (blue). Light shades around each plot presents standard error for each calculation. (C’) Probability Density Function (PDF) of RMSD sampled over the last 30ns (dashed line) of the simulations are shown in histograms.

ACE2 extracellular domain is a sizable receptor with about 600 amino acids; hence, it is expected that the overall RMSD of receptor would not change by the O-glycosylation of the RBD. As we aimed to study the alterations in ACE2-RBD interface, we calculated the RMSD of two *α*-Helices (Figures 7.B and 7.C) which are known as RBD interacting sites within ACE2. ^3^ Average RMSD plots of H1 and H2 show that the first helix is the most affected by the O-glycan (Figure 7.B and 7.C). H1 is more flexible in all O-glycosylated models (Figure 7.B). The histograms of the PDF for RMSD values show an increase in the peaks of models II and III for H1. H2 is less flexible in all the systems and is more persistent (Figure 7.C).

ACE2 RMSF plots support these observations, showing decreased fluctuations of H2 in the simulations (Figure 5.A). To highlight the differences, we have shown the relative differences compared to model 0 in Figure 5B.. The relative difference is presented as 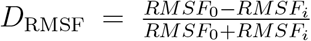 where *RMSF_i_* is per residue RMSF of each model. According to the definition more positive *D*_RMSF_ means lower relative RMSF.

The visualization of dynamics shows that O-glycan forms the most numerous polar contacts with H2 (five sites in H2 (N61,K68,A71,F72,E75) versus three sites in H1 (N38, L39, N49) Figure 8.) In contrast, according to the crystal structures, H1 forms more contacts with RBD than H2. ^3^RMSD plots of the attached O-glycans also show that the elongated models II and III are more flexible Figure 12. That is due to the several polar interactions that they form with H2. One should note that the RBD is not O-glycosylated in the Xray crystal structures. In fact that we see more interactions between O-glycan and H2 in our simulations suggests that the O-glycan keeps the RBD-ACE2 together by not interfering in the main ligand-receptor interactions takes place at H1.

**Figure 8:**
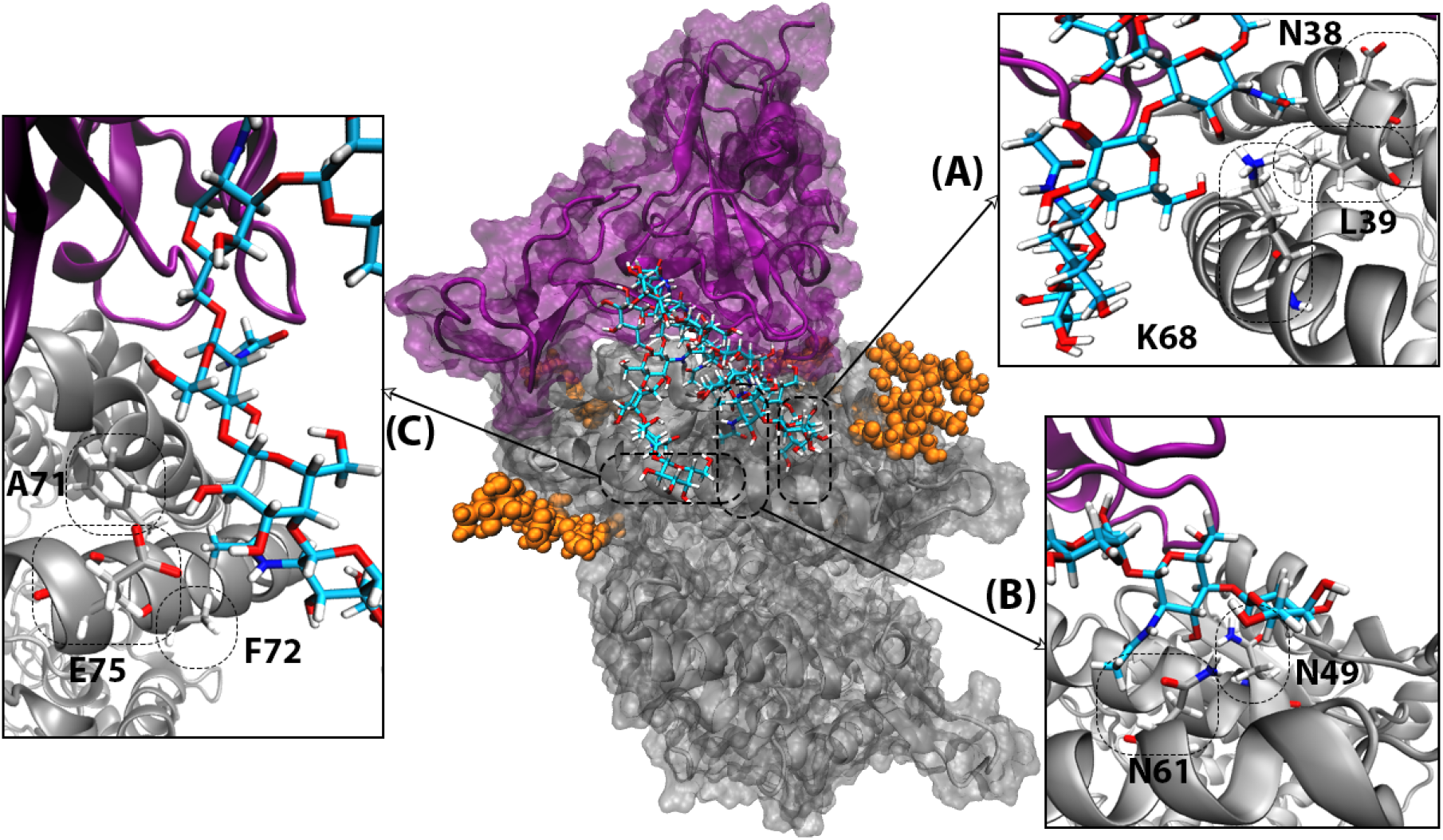
The extracellular domain of human ACE2 and RBD of the SARS-COV-2 S protein are shown with gray and purple respectively. The RBD attached O-glycan is shown with atom name sticks. N-glycans in the proximity of RBD-ACE2 interface are shown with orange spheres. Persistent polar interactions between H1 and H2 of ACE2 and the attached O-glycan are shown in A-C.

### Biding affinity of O-glycosylated RBD to ACE2

The decreased fluctuations of O-glycosylated models II and III, does not necessarily prove stronger RBD-ACE2 interaction. However, calculating the distance between RBD-ACE2, RBD-H1 and RBD-H2 could provide hints on this interaction. Distribution of the RBD-ACE2 distance values shows a similar pattern (peaks at 49.5 Å) in models I and III simulations (Figure 9.A). While, in model 0 simulations, RBD and ACE2 are by less than 1 Å closer (Figure 9.A). The distance fluctuation between RBD and ACE2 in model II is more altering.

**Figure 9:**
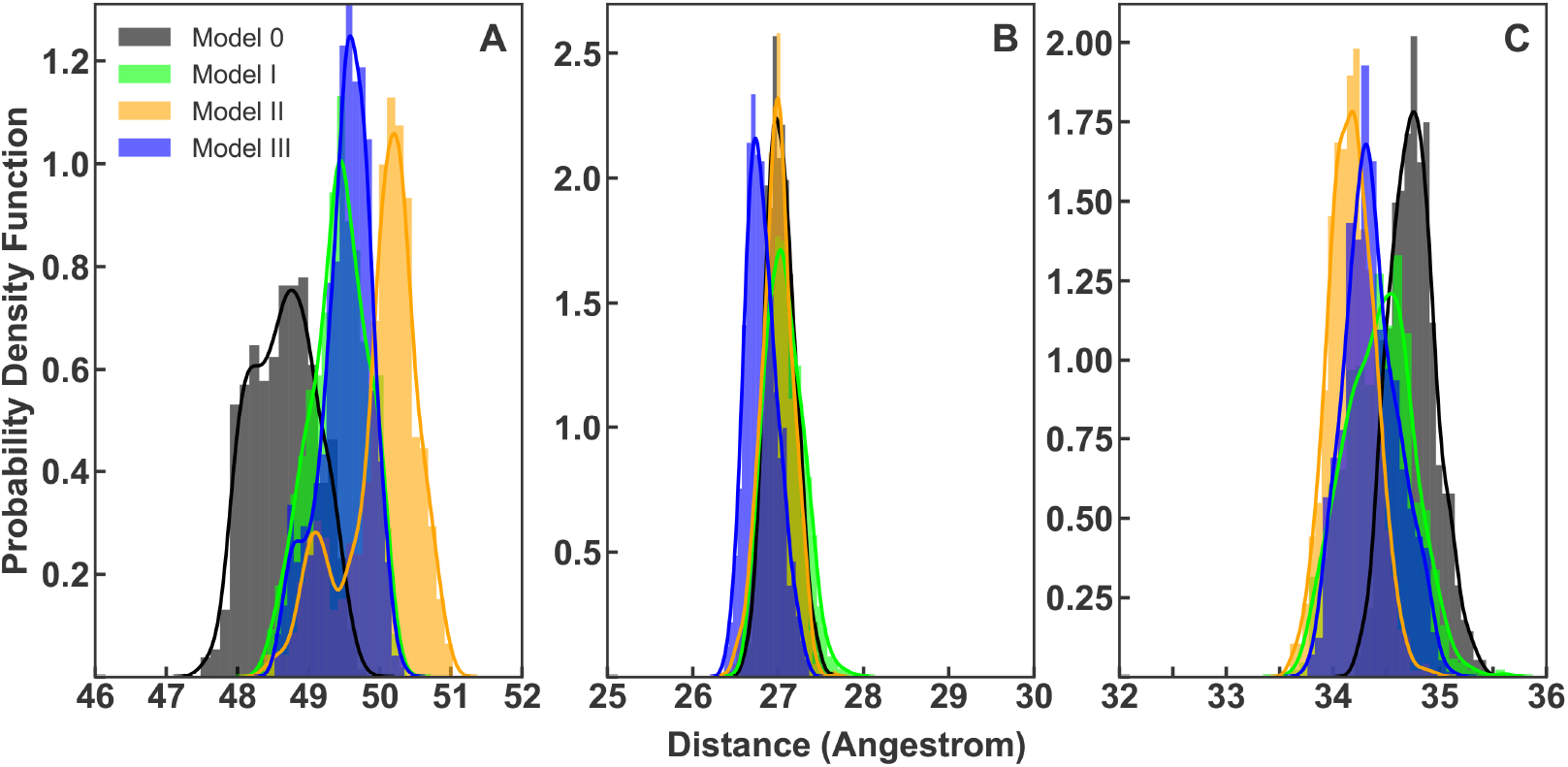
(A) PDF of RBD to ACE2 averaged distance distribution with a histogram and maximum likelihood gaussian distribution fit: calculated from all replicate of each system and are represented in black, lime, orange and blue for models 0-III respectively. (B) PDF of RBD to H1 averaged distance distribution with a histogram and maximum likelihood gaussian distribution fit: calculated from all simulations of each system and represented in black, lime, orange and blue for models 0-III respectively. (C) PDF of RBD to H2 averaged distance distribution with a histogram and maximum likelihood gaussian distribution fit: calculated from all simulations of each system and represented in black, lime, orange and blue for models 0-III respectively. Sampling were done from the last 30 ns of simulations.

In addition to the ACE2-RBD, distances between RBD, H1, and H2 of ACE2 were also measured (Figures 9.B and 9.C). Plots show that O-glycosylation always leads to a decrease between the RBD and the two helices (Figures 9.B and 9.C). This decrease is more significant in H2-RBD distance for all O-glycosylated models (Figure 9.C). While, H1-RBD distance distribution only presents a decrease in the most elongated attached O-glycan model (Figure 9.B). These observations suggest that RBD-ACE2 binding should be stronger at the direct interface upon O-glycosylation with oligosaccharides.

The binding free energy between the RBD and ACE2 also shows that the interaction is more favorable upon O-glycosylation in models II and III with longer attached glycans(Figure 10).

**Figure 10:**
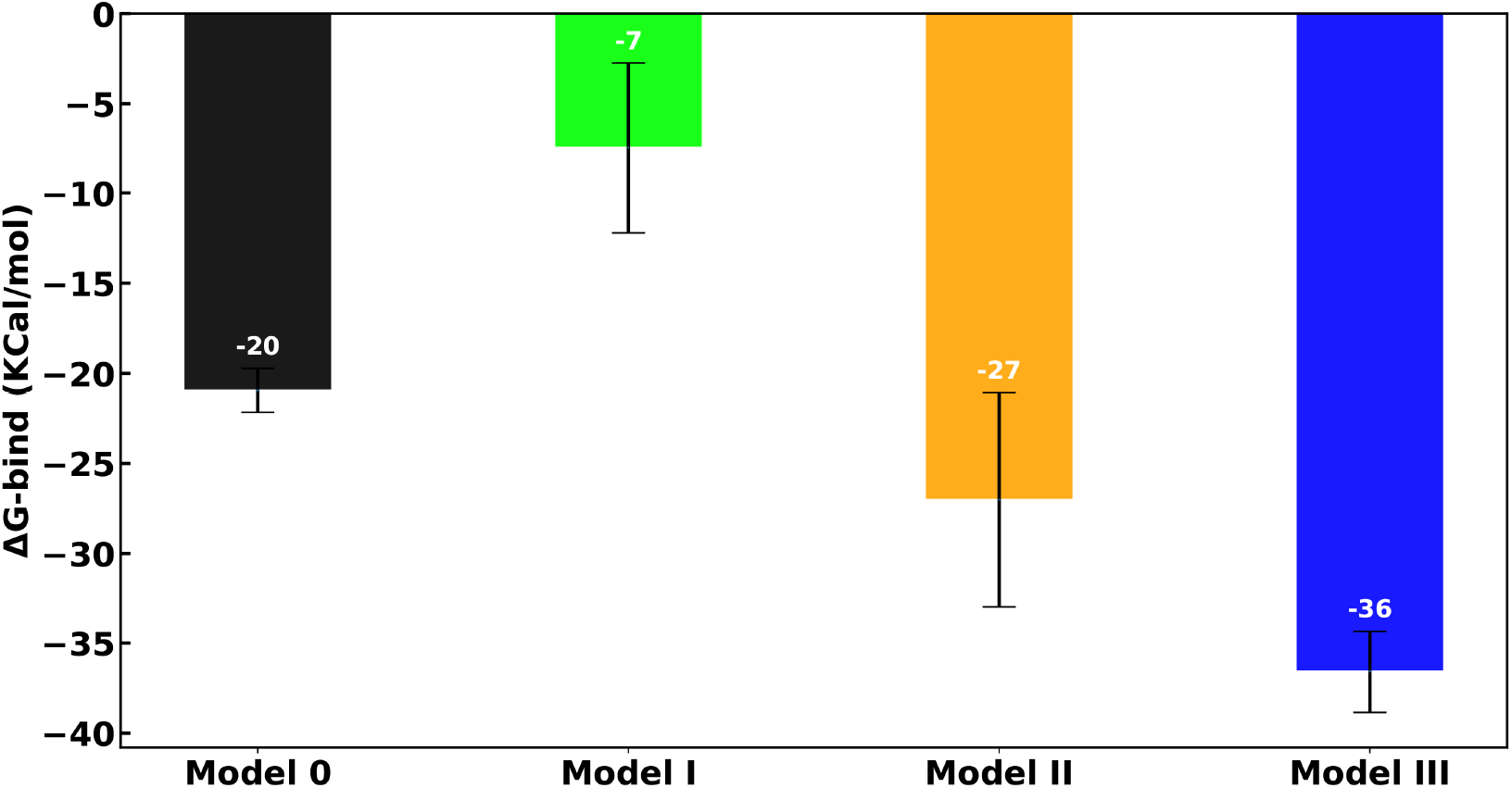
Relative binding free energy components for RBD-ACE2 complex. Overall binding free energy of models 0-III are shown in black, lime, orange and blue.

**Figure 11:**
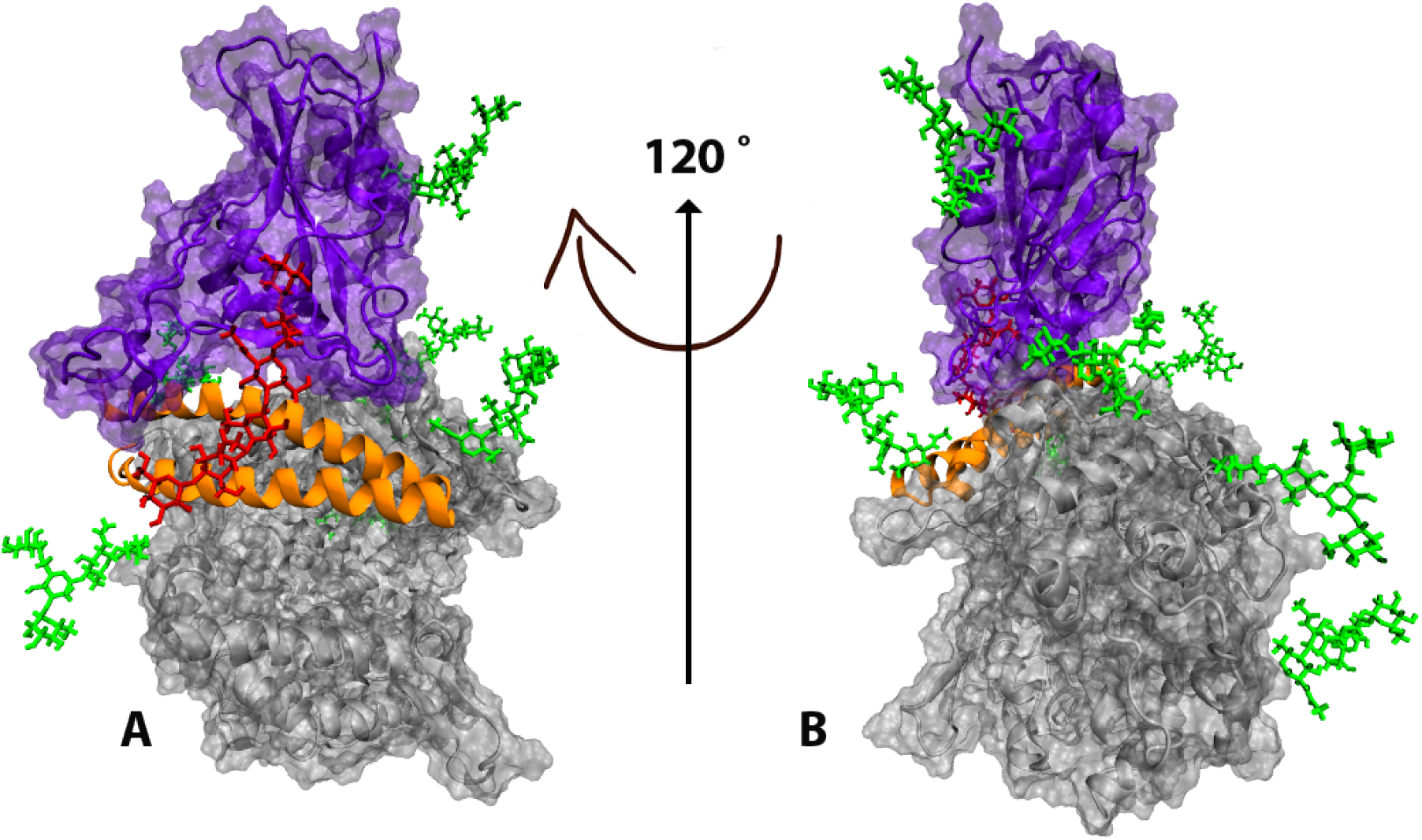
Visualization of the complex with full presentation of the N-glycans. 120° Z-axis rotation of model in A was presented in B. The extracellular domain of human ACE2 and RBD of SARS-COV-2 S protein is shown with gray and purple, respectively. The N-glycan are shown with green sticks. Red stick is model III o-glycan.

**Figure 12:**
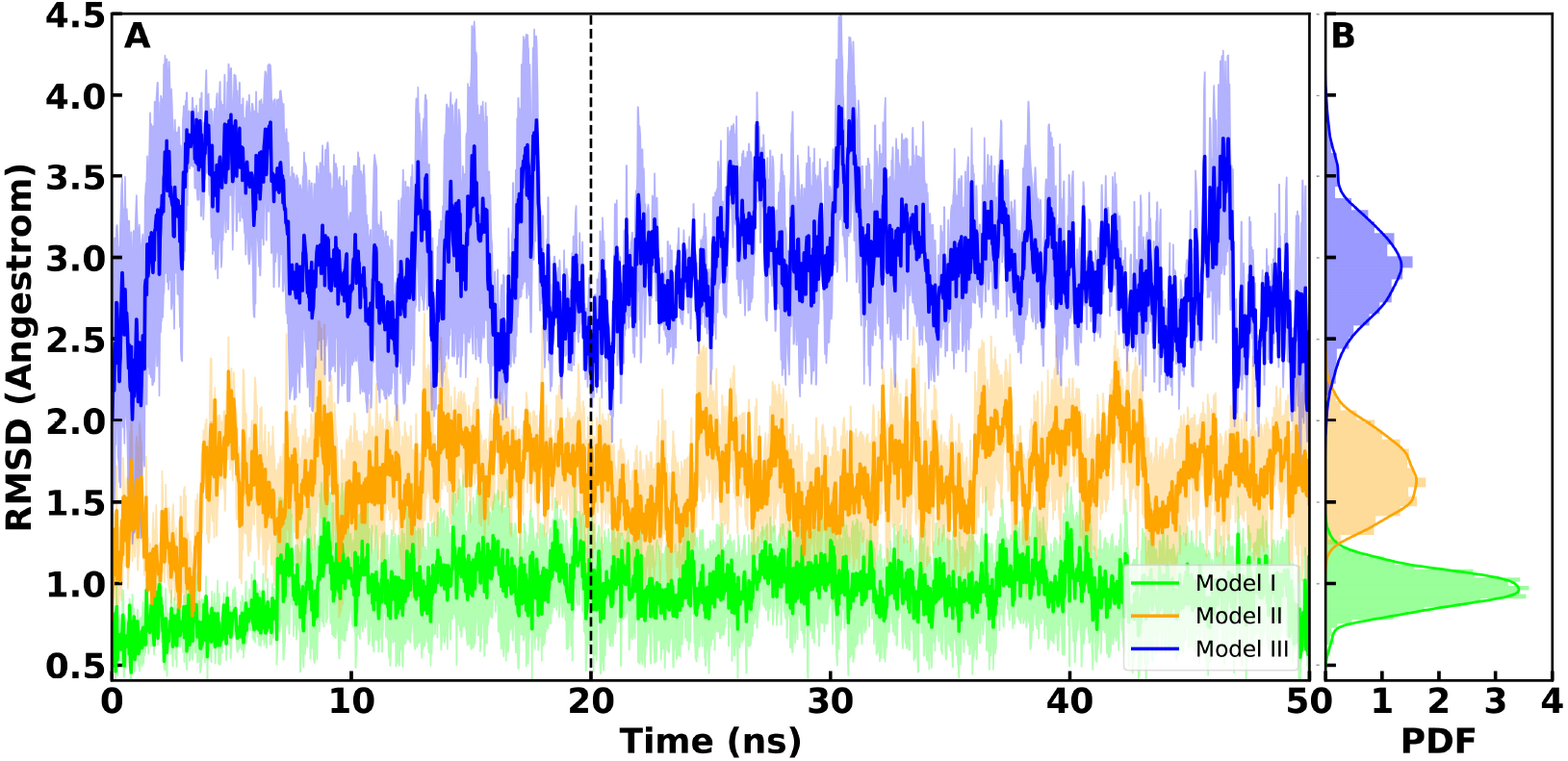
(A) Average RMSD plots for three models of O-glycans calculated from all replicates of each system and are shown for model I (lime), model II (orange) and model III (blue). Light shades around each plot presents standard error for each calculation. (B) Probability Density Function (PDF) of RMSD sampled over the last 30 ns (dashed line) of the simulations are shown in histograms

The binding free energies between RBD-ACE2, calculated with the MMPBSA method, show a monotonic decrease in ΔG by increasing the size of O-glycans (Figure 10) and staring from −20.94±1.22 kcal/mol for model 0 to −7.47±4.73 kcal/mol, −27.00±5.93 kcal/mol and −36.57±2.27 kcal/mol for models I, II and III, respectively (Figure 10). Interestingly, in model I with the core O-glycan the biding free energy is less favorable compared to model This might be due to the increased flexibility at the ACE2-RBD binding interface and the decrease of polar interactions made by the core glycan due to its small length. RMSD and RMSF plots of the RBD and overall ACE2-RBD (Figures 3, 6 and 4) are supportive of this observations and show an increase in the flexibility of the complex for model I. Chosen value of solute dielectric constant has shown to have a dramatic effect on the prediction of binding free energy, while using the MMPBSA method^39–41^ (see method section of Supporting information). Regardless of the given value to the dielectric constant, the decrease of binding free energy in the O-glycosylated models II and III with longer glycans is always persistent and statistically significant (Figure 10).

## Conclusions

S494 that only occurs in SARS-COV-2 and not in SARS-COV is located in direct RBD-ACE2 binding interface has the potential to be decorated by O-glycans. Previous experimental and computational studies emphasized the role of O-glycosylation in SARS-COV-2 high infectivity. The results of atomistic molecular dynamics simulations of SARS-COV-2 RBD in complex with ACE2 suggest that the O-glycosylation of S494 leads to the stronger interaction between RBD-ACE2 which can increase the virus infectivity. Three models of core and elongated O-glycans attached to RBD were tested here. We observed that attachment of the elongated O-glycans lead to less flexible ACE2-RBD dynamics and a decrease of distances between RBD and two major H1, H2 helices of ACE2 in the direct binding interface. Relative binding free energy of RBD-ACE2 is also more favorable in the O-glycosylated models with longer glycans. The increase of RBD binding affinity to ACE2 depends on the size of attached O-glycan. By increasing the size of O-glycan, the RBD-ACE2 binding affinity will increase. These findings add insightful information to the current status of SARS-COV-2-ACE2 glycosylation and its role in the virus’s high evasion rate. This hypothesis is a suitable target for experimental validation, and if proven to be critical, must be considered in further therapeutic designs.

## Acknowledgement

We are grateful to the HPC of SUT to provide the computational resources of this study. We are also thankful to H.Rajabi-Maham, H.Mobasheri, R.S.Haltiwanger and, R.J.Woods for their very useful suggestions.

## Supporting Information Available

### Supplementary Methods

For MMPBSA binding energy calculation ligand-receptor complex is first calculated in the gas state and then in solvated state. The overall relative binding free energy is the sum of those two ΔGs (Eq.1). Each ΔG can be decomposed into the contribution of different interactions and expressed in:^42^

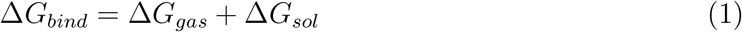

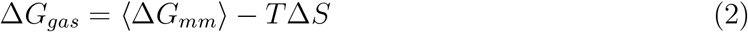

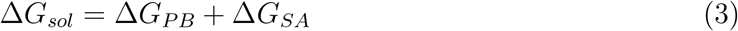

Where ⟨ΔG_*mm*_⟩, −TΔS, ΔG_*PB*_ and ΔG_*SA*_ are the ensemble-averaged protein-ligand interactions, entropy contributions, electrostatic, and nonpolar solvation energies, respectively. For calculating the entropic part of the binding energy, Interaction Entropy (IE) method^43^ was used:

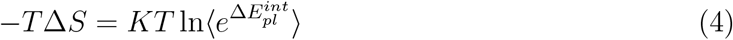

where 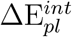 represents the fluctuation of protein-ligand interaction energy around the average energy and calculated as follow: ^43^

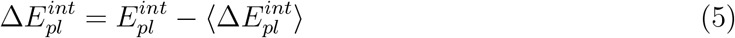

where 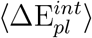 represents the average of protein-ligand interaction energy.

The 1.2 value for the dielectric constant in the binding free energy calculation was selected because it produced the closest values to the experimental RBD-ACE2 binding energy^2^ among all the other commonly used tested values. Also it has shown to be a reasonable value for such calculations according to the proposed fitting curve by Tingjun Hou et al.^40^

### Supplementary Tables

**Table 1:**
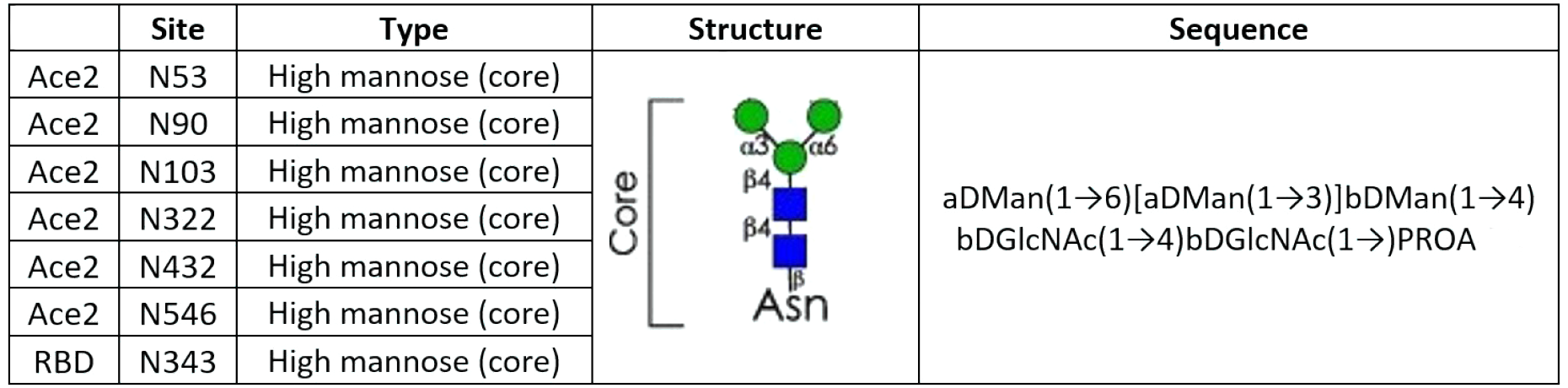
N-Glycan composition for ACE2 and RBD

---

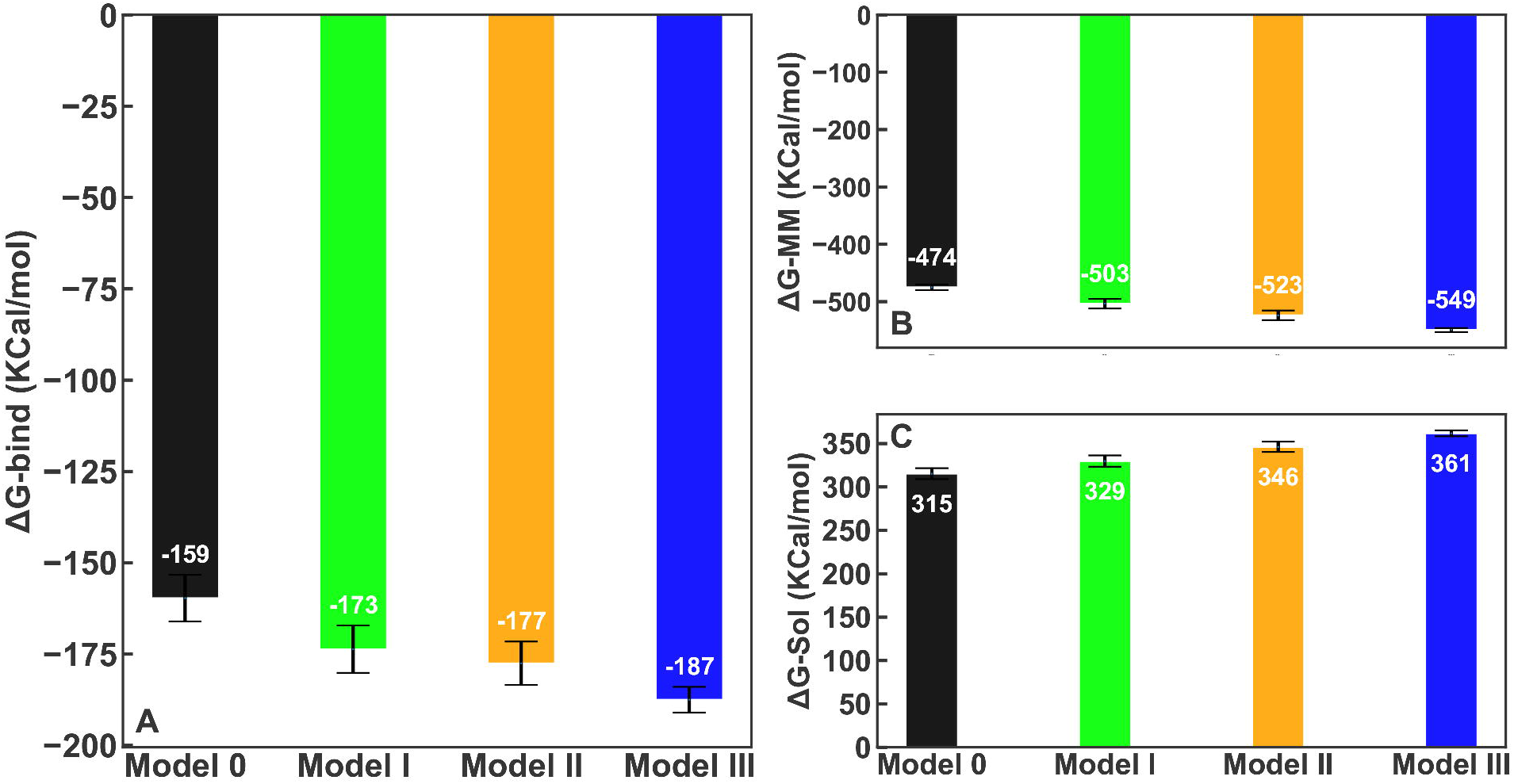

